# ESAT-6 of *Mycobacterium tuberculosis* downregulates cofilin1, leads to filamentous actin enrichment and reduces the phagosome acidification in infected macrophages, which are partially reversed by a single methionine mutation

**DOI:** 10.1101/2020.05.04.076976

**Authors:** P. P. Mahesh, R. J. Retnakumar, Sivakumar K C, Sathish Mundayoor

**Affiliations:** Mycobacteria Research, Bacterial and Parasite Disease Biology, Rajiv Gandhi Centre for Biotechnology, Thycaud P.O., Trivandrum 695014, Kerala, India; Genomics Core Facility, Rajiv Gandhi Centre for Biotechnology, Thycaud P.O., Trivandrum 695014, Kerala, India

## Abstract

*Mycobacterium tuberculosis* when phagocytosed by macrophages is not cleared completely and many of the bacteria remain in phagosomes indefinitely. In this study we considered abnormal retention of filamentous actin on early phagosomes contributing to defective phagosome acidification. The actin depolymerizing protein cofilin1 was found downregulated in macrophages infected with virulent *M. tuberculosis.* Also, phosphocofilin1, the inactive form of cofilin1, which leads to retention of filamentous actin, and the total filamentous actin itself were found upregulated in macrophages infected with virulent *M. tuberculosis*. Over expression of constitutively active cofilin1 in macrophages was found to decrease the level of filamentous actin and increase phagosome acidification when infected with virulent *M. tuberculosis*. The anticancer drug sorafenib which activates cofilin1 in PI3K dependent manner was also found to decrease the filamentous actin level and increase phagosome acidification. Cofilin1, known to be positively regulated by superoxide was found to be downregulated by ESAT-6 of *M. tuberculosis* where the latter is known to reduce ROS in macrophages. Ectopic expression of ESAT-6 in macrophages was found to downregulate cofilin1, increase filamentous actin and to transform the macrophages more spindle shaped. ESAT-6 was also found to decrease phagosome acidification in macrophages infected with an avirulent *M. tuberculosis* strain. Finally, this study proposes a role for the amino acid methionine in resisting ROS by creating M93 mutants of ESAT-6.

## 1. Introduction

Tuberculosis remains a global health concern and claims many lives every year, especially in Third World countries. The virulent *M. tuberculosis* after entry into macrophages, escapes phagosome-lysosome fusion and resides in the immature phagosome indefinitely[1]. The present study considers the role of filamentous actin in defective phagosome acidification. Actin filaments have a critical role in phagocytosis (formation of phagocytic cup), movement of endosomes and their fusion. But retention of excessive actin on phagosomes was reported to prevent phagosome-lysosome fusion[2]. The presence of actin on late phagosomes in virulent *M. tuberculosis* infection was shown to cause inadequate acquisition of the proton pump, V-ATPase by phagosome and inhibiting phagosome maturation[3].

Our study started when a differential expression of the actin depolymerizing protein, cofilin1, was observed in an *M. tuberculosis* infection experiment. Cofilin1 is one of the actin binding proteins having a pivotal role in actin dynamics[4,5]. Actin dynamics is necessary for diverse cellular functions including phagocytosis, movement of different endosomes and their fusion. Cofilin depolymerizes and severes filamentous actin (F-actin) which releases monomeric actin (G-actin) and also produces actin barbed ends which in turn serves for actin nucleation to form new filaments or creates actin branches. Cofilin1 is an 18kDa protein which is inactivated by Serine-3 phosphorylation. Serine-3 phosphorylated cofilin is not able to bind with actin. A coat of filamentous actin was found to prevent clustering of late endosomes and artificially targeting cofilin1 to late endosomes cleared the filamentous actin coat[6].

Cofilin1 is a stimulus responsive regulator of actin dynamics[7]. Dephosphorylation/activation of cofilin by various pro-inflammatory stimuli and hydrogen peroxide treatment were described earlier[8]. In a previous study we have mentioned that downregulating reactive oxygen species (ROS) may drive the macrophages to a more anti-inflammatory phenotype[9] and confirmed that an abundantly secreted protein of *M. tuberculosis* namely Early Secreted Antigenic Target-6kDa (ESAT-6) downregulates ROS in macrophages. The present study investigates the effect of ESAT-6 on cofilin1, filamentous actin and phagosome acidification. Finally, we propose a role for the amino acid methionine in resisting ROS by ESAT-6.

## 2. Materials and Methods Cell culture, bacteria and infection

THP1 monocytic cell line was maintained in RPMI-1640 medium (R4130, Sigma) supplemented with 10 % FBS. For obtaining macrophage monolayer THP1 cells with required cell density (3×10^6^cells in T-25 flask and 4×10^4^ cells per well in a 96 well plate) were treated with 20ng/ml of PMA (P8139, Sigma) and after one day PMA was washed off 2 times with RPMI. After washing the cells 3ml and 100µl complete medium was maintained until the addition of bacteria in T-25 flask and 96 well plate respectively (for 96 well plate bacteria were added in a100µl suspension). Stocks of *Mycobacterium tuberculosis* strains, H37Rv and H37Ra were prepared using beads (PROTECT bacterial preservers) and stored at-80^0^C. Periodically, the frozen stocks were revived in7H9 broth and streaked on LJ slants and maintained for at least 2 months in an incubator at 37^0^C. Prior to experiments bacteria were inoculated into 7H9 broth (271310, Difco) from LJ slants. 7-10 days old broth cultures were used for infection experiments. An OD of 0.15 at 610 nm was measured to give McFarland standard 1 and was calculated to contain 3×10^8^ bacteria/ml. On the day of infection, a log phase culture of bacteria was collected and passed through a syringe for 20 times and kept for 10 min to remove the clumps and the required volume of the culture was pelleted by centrifugation at 3000g for 10 min. Then the pellet was resuspended in RPMI and added to the macrophage monolayer. For increasing the infection efficiency, minimum volume of bacterial suspension was used to infect monolayer of macrophages. After the addition of bacteria, the macrophage monolayer was kept for 4h for phagocytosis. After 4h the extracellular bacteria were washed off 3 times with RPMI and complete medium was added.

### Chemicals

Chemicals and their respective concentrations used for the treatment of macrophages are listed below. H_2_O_2-_100µM, Sorafenib-5 µM (SML2653, Sigma), Wortmannin-100 nM (Sigma), Rifampicin-10µg/ml (Sigma)

### 2D gel electrophoresis

After 24h of infection, the total protein of infected and uninfected macrophages was isolated using 2D lysis buffer composed of 8M urea, 1M thiourea and 4% CHAPS along with 1mM PMSF. After 30 minutes of incubation at room temperature the cell lysate was centrifuged at 14000g for 20 min at 18^0^C and then 300µg of total protein was loaded on a 7cm IPG strip of 3-10 pI range (163-2000, BIO RAD). After isoelectric focusing, the strips were equilibrated, and PAGE (15% gel) was run to resolve the proteins. From the gel, a visually identified spot was picked and analyzed by MALDI-TOF-TOF.

### Quantitative RT-PCR

After 12h of infection, total RNA of infected and uninfected macrophages was isolated using Illustra RNAspin (25-0500-71, GE) and the concentration was estimated by NanoVue Plus, GE. Primers for the genes were designed by Primer Premier Software. Primer pair for the reference gene GAPDH: FP: 5’ TCAAGAAGGTGGTGAAGCA 3’, RP: 5’ AGGTGGAGGAGTGGGTGT 3’. Primer pair for cofilin: FP: 5’GCCTGAGTGAGGACAA3’, RP: 5’ GACAAAGGTGGCGTAG 3’.Primer efficiency for cofilin: 93.4% and GAPDH: 92.15%. qRT-PCR was performed using iScript one step RT-PCR kit (170-8892, BIO RAD). The relative quantitation and calculation of primer efficiency were done by the same software (CFX manager, BIO RAD) which performed the RT-PCR

### Western blot

At various time points, infected and uninfected macrophage monolayers were washed with ice cold PBS, scraped in 1ml PBS and were centrifuged at 2000 rpm for 5 min at 4^0^C. The cell pellet was lysed with RIPA buffer (R0278, Sigma) in the presence of protease inhibitor cocktail (P8340, Sigma). After 5min of incubation on ice, the cell lysate was centrifuged at 14000g for 10min at 4^0^C and supernatant was collected. Protein was estimated using a BCA kit (23225, Thermo Scientific) and loaded at 15µg per well. β-tubulin (ab6046, abcam) was used as loading control. Blot was probed by anti-cofilin antibody (ab42824, abcam) or anti-cofilin –phospho S3-antibody (ab131274, abcam), and subsequently with anti-rabbit secondary antibody HRP (sc-2004, Santa Cruz) or anti-mouse secondary antibody (Sigma), and was developed using ECL Prime (RPN2232, GE) or using manually prepared luminol (Sigma). Relative expression was calculated by taking the ratio of intensities of respective proteins to that of β-tubulin using Quantity One or Image lab software (BIO RAD) Figures were processed and presented using Adobe Photoshop-7 and Microsoft Power Point.

### Confocal microscopy

THP1 cells were seeded in an optical bottom 96 well plate at 4×10^4^ cells per well and induced to form macrophage monolayer. The cells were fixed by 4% paraformaldehyde, permeabilized by 0.1% Triton X100 and blocked with 3% BSA and 0.3M glycine in PBS. Images were taken under 40X objective with a confocal microscope (Nikon).

Phalloidine Red staining: After fixation and permeabilization of macrophage monolayers incubation of a working solution of the dye (Acti-stain 555, PHDH1, Cytoskeleton) in PBS at 100nM (stock-14µM in methanol) was done at room temperature in dark for 30min. Then, washed 3 times with PBS and imaged.

Lysotracker Red staining: a 100nM concentration of the dye (Invitrogen) was prepared in RPMI and incubated the cells for 40min in a CO_2_ incubator. Then, washed 3 times with PBS and processed for imaging by confocal microscopy.

Figures were processed and presented using Adobe Photoshop-7

### DCFDA fluorescence measurement by flow cytometry

Macrophages were seeded at 3 million/T-25 flask, washed 3 times with RPMI, detached the cells using a non-trypsin reagent (Sigma) and suspended the cells in RPMI containing 10µM DCFDA, incubated in CO_2_ incubator for 30min, centrifuged, washed twice with PBS and analyzed by flow cytometry using 488nm laser.

### Plating and colony count

To find out the effect of sorafenib treatment on the survival of bacteria, macrophages were infected with H37Rv. After 4h of phagocytosis gentamicin was added to the medium at 15µg/ml and incubated for 1h to kill the extracellular bacteria, then washed 3 times with RPMI. Sorafenib was treated after one day of infection, continued the treatment for another two days and after that the cells were lysed with sterile distilled water, diluted to 10^-2^ and plated on 7H10 agar (262710, Difco) supplemented with 0.1% casitone (0259-17, Difco). Colonies were counted after 3 weeks.

### Ectopic expression of ESAT-6 in macrophages

ESAT-6 was amplified from H37Rv DNA using the following primers: FP: 5’ CG<u>GAATTC</u>CGATGACAGAGCAGCAGTGGAAT3’ with EcoR1 site and RP: 5’CG<u>ACGCGT</u>CGCTATGCGAACATCCCAGTGACG3’ with Mlu1 site. The amplified fragment was cloned into MCS A of pIRES (Clontech). The reporter gene ECFP was taken from pECFP-N1 (Clontech) by restriction digestion at Sal1 and Not1 sites and ligated into MCS B of pIRES. The insertions were confirmed by both restriction digestion and sequencing. The plasmid was isolated using an endotoxin free plasmid isolation kit (EndoFree Plasmid Maxi Kit, Cat. No. 12362, QIAGEN). Before transfection THP1 cells were washed in serum free RPMI for 2 times. Then a 10-20 µg of plasmid was mixed with 40×10^6^ THP1 monocytes suspended in serum free RPMI in a 4mm electroporation cuvette and electroporated at the given conditions; choose mode-LV, pulse length-5ms, charging voltage-350V in a BTX Harvard Apparatus, ECM 830[10]. Cells were incubated for 10min on ice before and after electroporation and transferred to culture flasks containing RPMI+10%FBS and incubated for 24hrs. After 24hrs the viable cells were counted using Trypan blue (T8154, Sigma) staining, seeded into plates at respective densities and treated with PMA for next 24hrs.For Western blot a 10µg of total protein was loaded. Transfection of cells was confirmed by imaging the cells with 405nm laser for ECFP expression with a confocal microscope Creation of ESAT-6 M93 mutants: M93A and M93C mutants of ESAT-6 were made by manually changing the codon for methionine to alanine or cysteine in the reverse primer of ESAT-6 gene and cloning was done as described earlier.

*E. coli* containing pIRES plasmid with ESAT-6 or its mutants were used to check ROS resistance since the immediate early CMV promoter in the plasmid is active in the bacteria also[11].

### F-actin/G-actin ratio

F-actin stabilization buffer: 0.1M PIPES at pH-6.9, 30% glycerol, 5% DMSO, 1mM MgSO_4_, 1mM EGTA, 1% IgePal and 1mM ATP F-actin depolymerization buffer: 0.1M PIPES at pH-6.9, 1mM MgSO_4_, 10mM CaCl_2_ and 8M urea. The cells were lysed with actin stabilization buffer on ice for 10min and centrifuged at 4^0^C for 75min at 16000g. Supernatant was collected, containing the G-actin portion. The pellet containing F-actin was solubilized in the depolymerization buffer. Both the F-actin and G-actin portions were loaded at 10µl each with loading dye. The Western blot was probed with an anti-β-actin antibody (Sigma).

### MTT assay

MTT reagent (M2128, Sigma) was dissolved in PBS at 5mg/ml. 20μl of MTT solution was added to 200μl of culture medium of infected macrophages in each well of a 96 well plate and incubated for 4h. After the incubation the solution was aspirated and 100μl of DMSO was added to each well. The content was mixed by keeping the plate on a rocker for 5 min and absorbance was measured at 570 nm. Background absorbance was measured at 690 nm and subtracted from the value at 570 nm.

### Statistical analysis

Statistical analysis was done using IBM SPSS statistics software, version 19. Comparison of means was done by one way ANOVA (Tukey’s method) by setting alpha value as 0.05. Values were expressed as mean ± SE.

## 3. Results

### 3.1. *Mycobacterium tuberculosis* downregulates cofilin1 in macrophages

Macrophage monolayers were infected with either live virulent *M. tuberculosis*, H37Rv or heat-killed H37Rv at MOI-1:20 and uninfected macrophages were used as control. Total protein was isolated 24h post infection and resolved by 2-dimensional gel electrophoresis (2DE) (Fig. 1A). One of the proteins, which was downregulated in live H37Rv-infected macrophages when compared to heat-killed H37Rv-infected macrophages was identified as cofilin1(non-muscle isoform) by MALDI-TOF-TOF. To confirm the differential expression of cofilin1, quantitative real time PCR (RT-PCR) and Western blot (WB) were done (Fig. 1B & 1C) and results like that in 2DE were obtained. Also, WB showed that the expression of cofilin1 in the avirulent *M. tuberculosis* strain H37Ra-infected macrophages was like the expression in heat-killed H37Rv-infected macrophages (Fig. 1C). Further, phosphorylation of cofilin1 in macrophage was checked by WB under the infection conditions described earlier using an antibody specific to Serine-3 phosphorylated cofilin1. Phosphorylation of cofilin1 was found to be upregulated in live H37Rv-infected macrophages compared to both heat-killed H37Rv-infected and H37Ra-infected macrophages (Fig. 1C). The above results show that live virulent *M. tuberculosis* downregulates cofilin1 expression and inactivates it by increasing its phosphorylation in macrophages compared to non-pathogenic infection conditions. While a significant change in expression of cofilin1 was seen around 24h post infection, a significant difference in phosphorylation was visible at zero hour of infection (4h after addition of bacteria to macrophages) (Fig. 1D).

**Figure 1.**
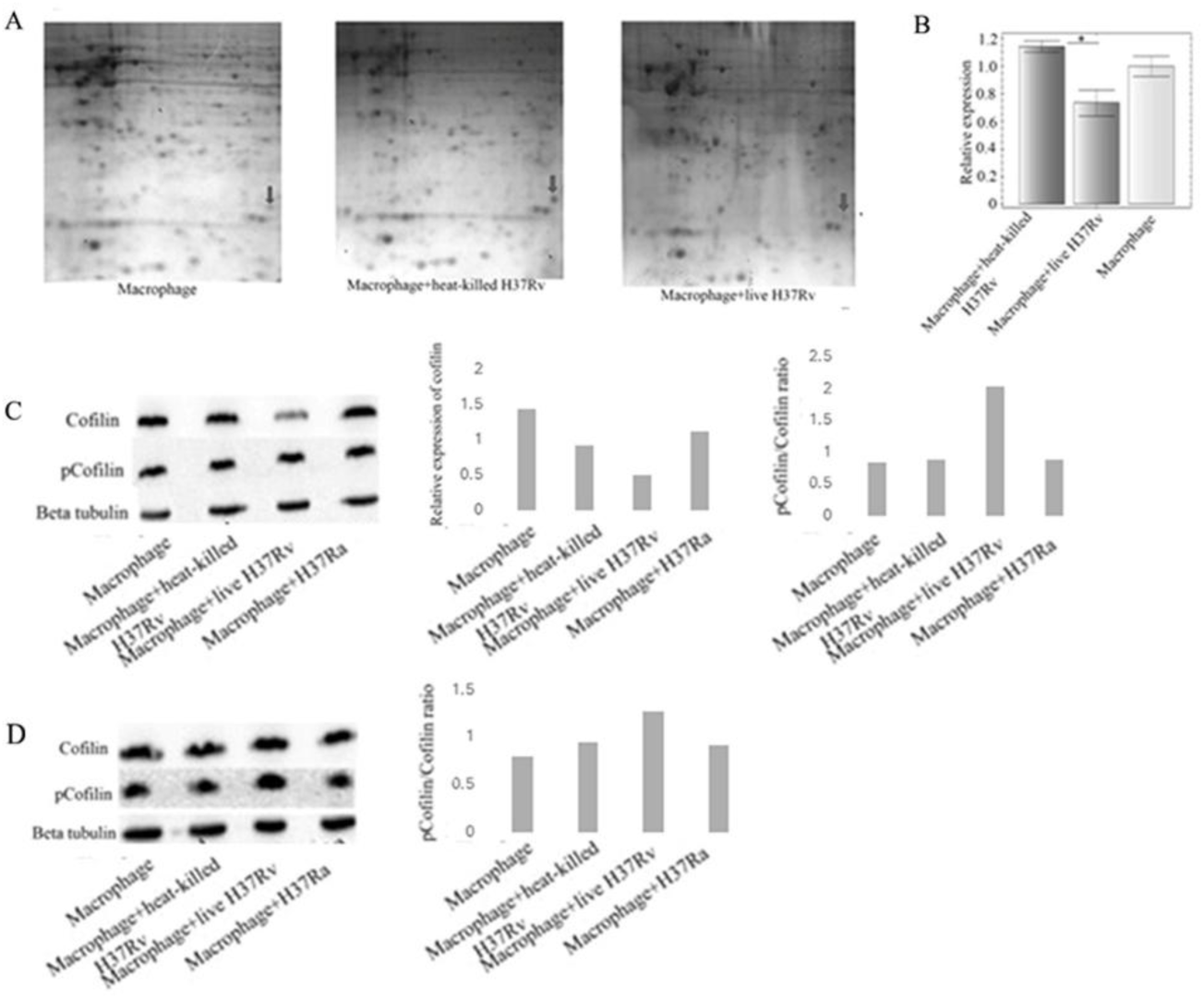
*M. tuberculosis* downregulates cofilin1. A) 2DE images showing the spot for cofilin in different infection conditions. B) RT-PCR showing differential expression of cofilin gene at 12h post infection. Values are expressed as mean±SE. n=3. * *p*<0.05. C) Differential expression of cofilin and pCofilin in different infection conditions at 24h post infection. D) Differential phosphorylation of cofilin at zero hour (4h after addition of bacteria to macrophages) in different infection conditions.

### 3.2 Connecting Cofilin to phagosome acidification/phagosome-lysosome fusion

Since cofilin1 is an actin depolymerizing protein and an abnormal retention of filamentous actin on early phagosomes is known to reduce phagosome acidification, we first investigated the filamentous actin content of macrophages in different infection conditions. We could observe an increase in F-actin/G-actin ratio (polymerized versus depolymerized actin) in live H37Rv-infected macrophages compared to heat-killed H37Rv-infected and H37Ra-infected macrophages (Fig.2).

**Figure 2.**
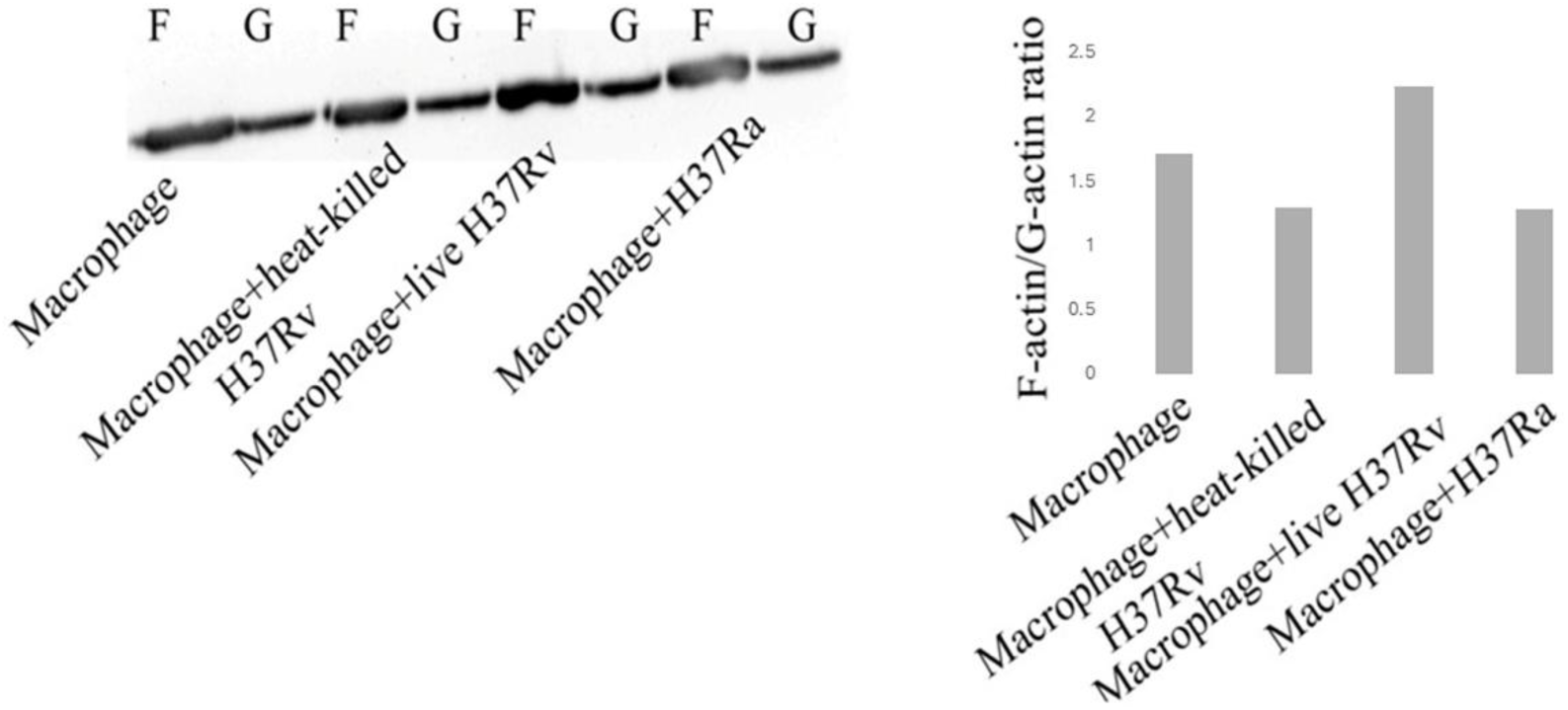
F-actin content of macrophages is increased on virulent *M. tuberculosis* infection. Figure shows F-actin/G-actin ratios in different infection conditions. The results are representative of two independent experiments, Cofilin1 was reported to be dephosphorylated by the broad spectrum kinase inhibitor and anticancer drug sorafenib[13] and the mechanism of this dephosphorylation/activation of cofilin1 was found to be through Phosphatidylinositol-3-kinase (PI3K) dependent activation of cofilin1 phosphatase SSH (Sling Shot Homologue). The above study also showed a thinned F-actin network in sorafenib-treated cells. We treated the macrophages with 5µM sorafenib after phagocytosis of bacteria (4h after addition of bacteria) and found that F-actin mesh appeared to be thinned in H37Rv-infected macrophages which otherwise showed rich F-actin on visual examination (Fig. 3A).

It is worthy to connect this observation to an earlier study which showed slower movement of lysosomes in virulent *M. tuberculosis* - infected macrophages compared to non-pathogenic mycobacteria-infected macrophages[12].

Further, the phagosome acidification greatly increased in sorafenib-treated H37Rv-infected macrophages (Fig. 3B). These effects of sorafenib were reversed when the sorafenib-treated macrophages were treated along with the PI3K inhibitor wortmannin.

**Figure 3.**
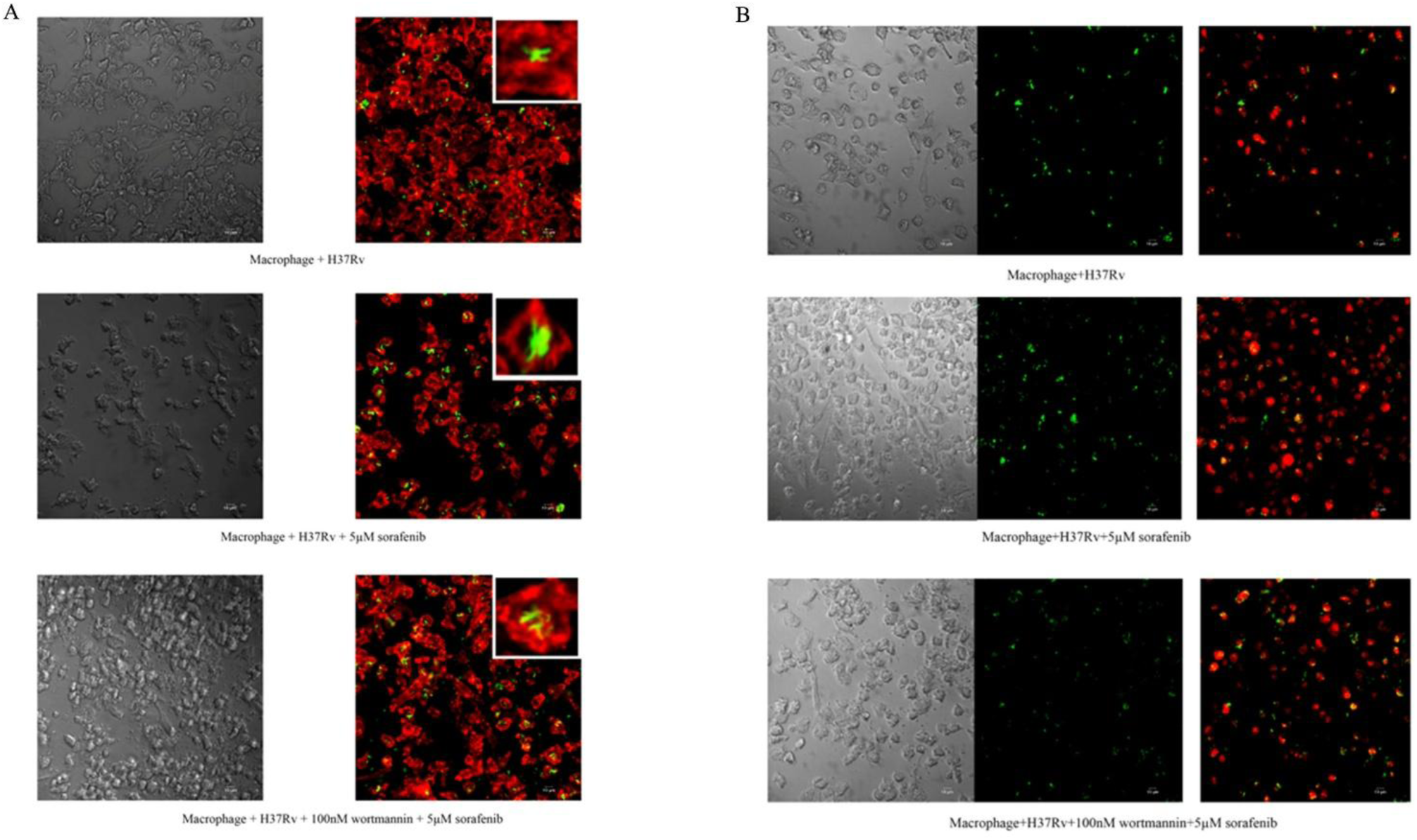
A) Sorafenib treatment reduces the F-actin content (as visualized by Phalloidine red) of *M. tuberculosis* (GFP expressing, green)-infected macrophages and PI3K inhibitor wortmannin reverses the effect of sorafenib. **B)** Sorafenib treatment increases phagosome acidification (as visualized by Lysotracker red) of virulent *M.tuberculosis* (GFP expressing, green)-infected macrophages and PI3K inhibitor wortmannin reverses the effect of sorafenib. Images are representatives of three independent experiments.

We also obtained the colony count of H37Rv-infected macrophages treated with 5µM sorafenib. In this case sorafenib was added one day after infection to allow the establishment of infection by virulent bacteria and sorafenib treatment was continued for another two days. The colony count from sorafenib-treated macrophages was significantly lower than that from the untreated control (Fig. 4A). Since sorafenib is a pro-apoptotic drug, cell death was noticed in macrophages treated with 1µM and 5µM concentrations of sorafenib compared to untreated macrophages, as calculated by MTT assay (Fig. 4B). 7.5 µM and 10µM concentrations of sorafenib drastically increased the cell death (data not shown).

**Figure 4.**
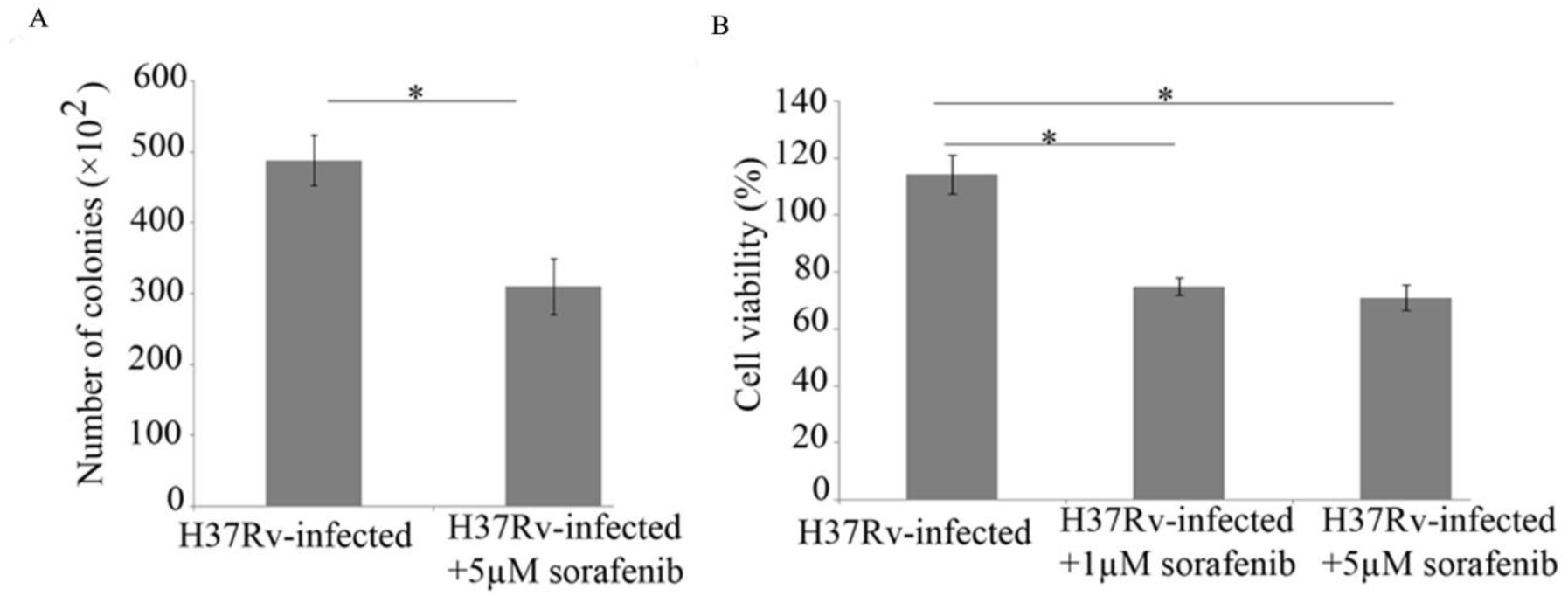
Sorafenib treatment of infected macrophages kills *M.tuberculosis*. **A)** Colonies of H37Rv from infected macrophages with or without sorafenib treatment, plated after diluting to 10^-2^. N=10, *p*<0.05. **B)** MTT assay of macrophages treated with 1µM and 5µM sorafenib. N=6, *p*<0.05. Cell viability in all cases was compared to uninfected macrophages. Values are expressed as mean±SE.

To further strengthen the above observations, we constructed a pIRES plasmid bearing constitutively active cofilin1-S3A mutant (plasmid construct from Addgene[14]) with ECFP reporter downstream of the gene. Since Serine-3 phoshorylation inactivates cofilin1, the mutation from serine to alanine turns cofilin1 readily active. The macrophages transfected with cofilin1 mutant or plasmid control were infected with H37Rv, and both F-actin distribution and phagosome acidification were checked 24h post infection. It appeared to be a significant thinning of F-actin mesh on visual examination (Fig. 5A) and increased phagosome acidification (Fig. 5B) in cofilin1 mutant plasmid-transfected macrophages compared to plasmid control-transfected macrophages.

**Figure 5.**
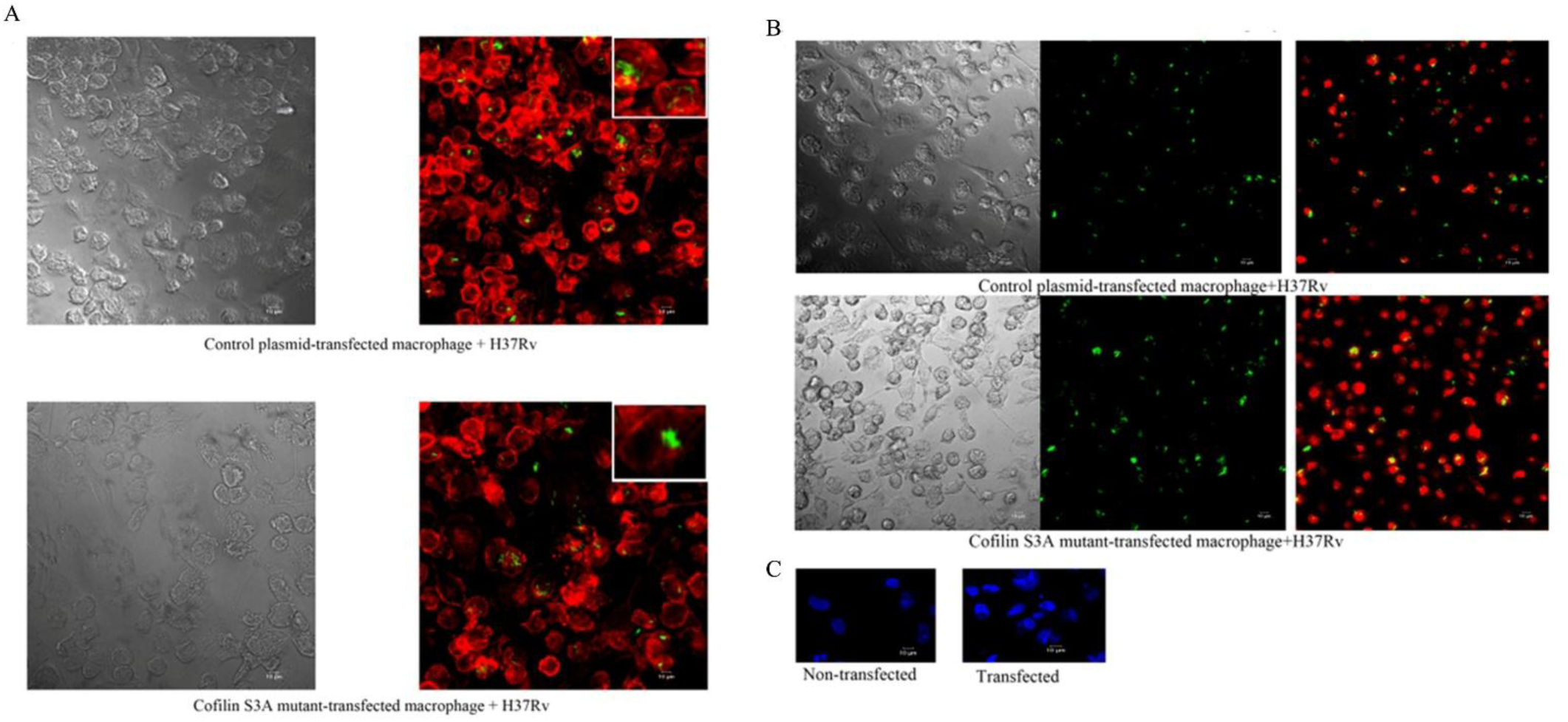
Transfection of S3A mutant of cofilin1 decreases F-actin content and increases phagosome acidification in virulent *M. tuberculosis*-infected macrophages. **A)** Phalloidine red staining of H37Rv (GFP expressing, green)-infected macrophages overexpressing constitutively active cofilin. **B)** Lysotracker Red staining of H37Rv (GFP expressing, green)-infected macrophages overexpressing constitutively active cofilin. **C)** Spreading of blue color in transfected macrophages shows the ECFP expression compared to non-transfected macrophages with the blue color confined to nucleus alone, when DAPI-treated macrophages were imaged with a 405nm laser. Images are representatives of three independent experiments.

### 3.3 Effect of ESAT-6 of *Mycobacterium tuberculosis* on actin and cofilin1: proposing a mechanism of action for ESAT-6

A previous study showed that *M.tuberculosis* ESX secretory system and its effectors cause the arrest of phagosome maturation in infected macrophages[15]. Another study showed that ectopic expression of ESAT-6 in *Dictyostelium* induces the formation of an F-actin coat around *M.tuberculosis* phagosome, and it helps them to escape from phagosome-lysosome fusion and spread to uninfected host cells by non-lytic ejection[16]. So far in this study we have correlated an increase in F-actin in *M.tuberculosis*-infected macrophages to reduced phagosome acidification. Now we propose a mechanism of action of ESAT-6 which might have resulted in the above observations.

In a previous study we presented our findings which support the assumption that downregulation of reactive oxygen species (ROS) may drive the macrophages to a more anti-inflammatory phenotype[9]. A balanced inflammation is needed for the macrophage to efficiently contain the invading microbes[17]. It was reported that tuberculous granuloma contains more M2 macrophages (anti-inflammatory phenotype) than M1 macrophages (pro-inflammatory phenotype)[18]. We assume that secretory products of *M.tuberculosis* induce this anti-inflammatory effect. Also, ESAT-6 and CFP-10 were reported to downregulate ROS in macrophages[19]. In this context it is noteworthy that chronic granulomatous disease (CGD) which is characterized by severe, protracted and often fatal infection, results from a failure of the NADPH oxidase enzyme system in the patient’s phagocytes to produce superoxide[20].

To this end we hypothesize that a direct scavenging of ROS by ESAT-6 is possible and propose that the amino acid methionine in the exposed portions of the protein may react with superoxide/ROS to form methionine sulfoxide that in turn reduces the ROS concentration needed to drive effective pro-inflammatory signaling. Also, the cytoplasmic methionine sulfoxide reductase of macrophage may convert the methionine sulfoxide formed in ESAT-6 back to methionine and expose it again to ROS. The antioxidant function of methionine is well described in the literature[21,22] and it is more sensitive to ROS than cysteine in protecting active sites of various enzymes[23]. The disordered regions of Prion proteins which extend out of neuronal plasma membranes contain multiple numbers of methionine and they scavenge oxidants[24]. It is interesting to note that a study of the structure of ESAT-6.CFP10 complex shows that it does not have pore forming, nucleic acid interacting and catalytic properties[25], which gives support to our assumption.

There are 3 methionines in ESAT-6 including the starting one. We selected M93 for mutation to find its effect on ROS resistance. M93 is present in the C-terminal flexible arm of ESAT-6 which does not contribute to the structure of the protein. At the same time mutations in C-terminal flexible arm including the mutation of methionine were reported to cause the loss of virulence imparted by ESAT-6[26].

Methionine 93 was mutated to alanine or cysteine. ESAT-6, ESAT-6 M93A and ESAT-6 M93C expressing *E. coli* were treated with 400µM H_2_O_2_ in broth culture and incubated for 6hrs and optical densities (OD_610 nm_) were measured in every hour. The OD of *E.coli* containing the control plasmid remained zero throughout the period. *E. coli* containing ESAT-6 had the highest OD at all time-points followed by E. *coli* with ESAT-6 M93A and ESAT-6 M93C respectively (Fig. 6A). After 6hrs, ODs of all broth cultures became similar (data not shown), probably due to the degradation of H_2_O_2_. Even though the mutation of methionine to cysteine was designed to rescue the antioxidant function imparted by the presence of sulfhydryl groups in both amino acids, the effect of M93C mutant was nearer to the control in the above experiment. This may be due to the formation of double-sulfide bond between cysteines of individual protein units to cause the dimerization of ESAT-6. There is no cysteine in ESAT-6 or CFP-10 and many of the secreted virulent proteins of different bacteria do not contain cysteine[27].

**Figure 6.**
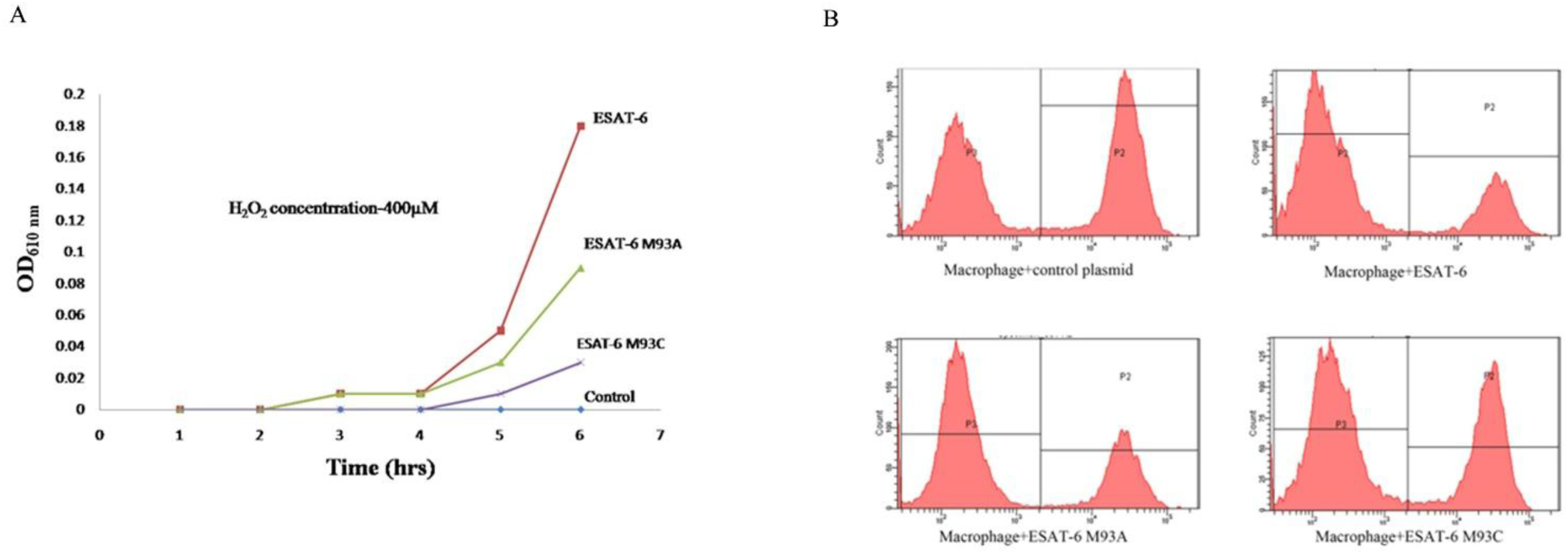
ESAT-6 makes *E.coli* resistant to hydrogen peroxide treatment and reduces the level of ROS in macrophages. **A)** ESAT-6 and its mutants-expressing *E. coli* were grown in broth culture treated with 400µM H_2_O_2_ and ODs were measured. **B)** ESAT-6 downregulates ROS (as measured by DCFDA fluorescence by flow cytometry) and M93 mutation partially reverses this effect in transfected macrophages 24h post PMA induction. The images are representatives of 3 independent experiments.

Further, we transfected ESAT-6 and its mutants in macrophages and measured ROS after 24h post PMA induction using DCFDA fluorescence as a readout by flow cytometry. The result was similar to what was obtained from the *E.coli* experiment (Fig. 6B). The graphs for ESAT-6 and ESAT-6 M93A mutant look similar, but the area of low fluorescence populations is more in the case of ESAT-6 (and the high fluorescence population is less to some extent) compared to the mutant while ESAT-6 M93C was closer to control.

Cofilin1 expression was checked in ESAT-6 and its mutants-transfected macrophages at 24h post PMA induction by WB. Figure 7 shows reduced expression of cofilin1 in ESAT-6-transfected macrophages compared to plasmid control-transfected macrophages. Macrophages transfected with ESAT-6 mutants and ESAT-6-transfected macrophages treated with 100µM H_2_O_2_ showed increased expression of cofilin1 compared to ESAT-6-transfected macrophages.

**Figure 7.**
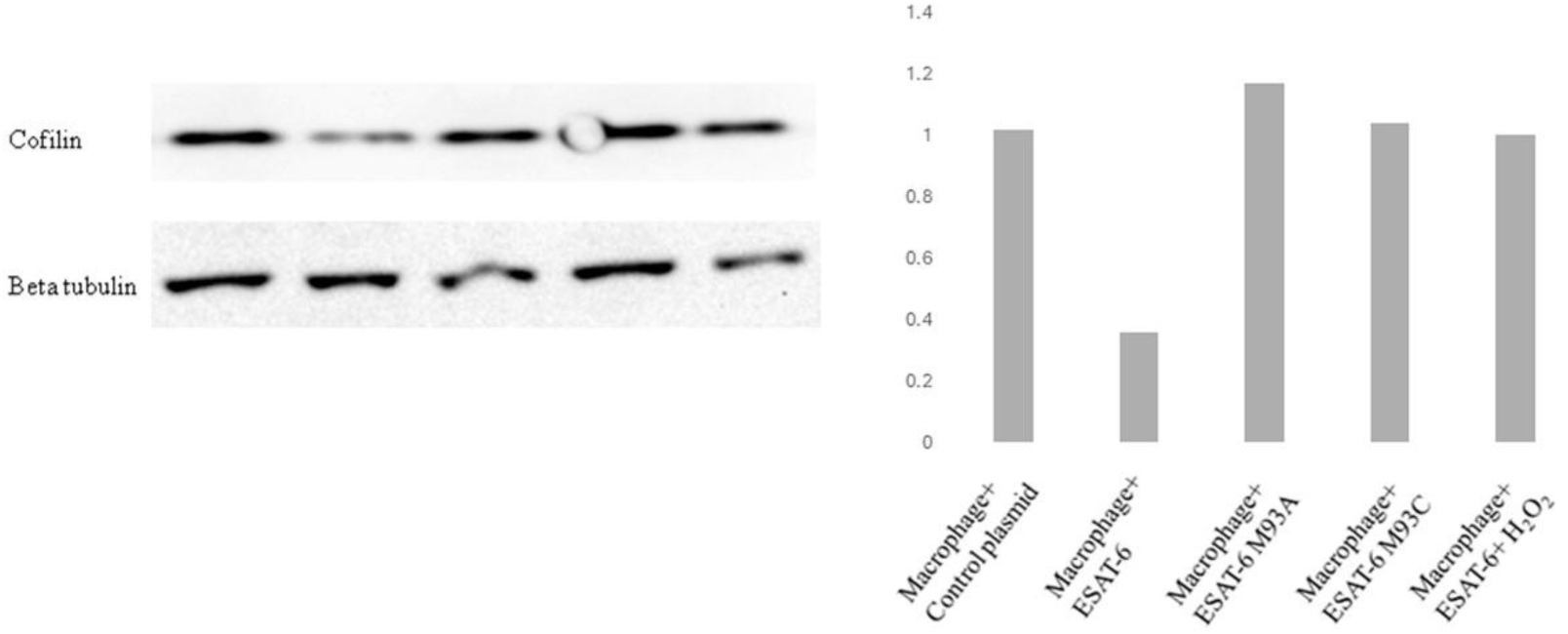
ESAT-6 downregulates cofilin1 and this effect is reversed by its M93 mutation and H_2_O_2_ treatment in transfected macrophages. Figure shows the WB done in macrophages after 24h of PMA induction.

We also checked the effect of transfection of ESAT-6 and the mutants on F-actin and phagosome acidification. We got more spindle shaped (M2 like) macrophages which appeared to have enriched F-actin network in cytoplasm on visual examination, in the case of ESAT-6 transfection (Fig. 8A). Phagosome acidification was also found to be reduced in ESAT-6-transfected macrophages (Fig. 8B). These effects were partially nullified in the case of macrophages where ESAT-6 mutants were transfected. In this experiment, rifampicin-treated (10µg/ml in infection medium) H37Ra was used for infection to ensure that no protein from the bacterium interfered in the infection process.

**Figure 8.**
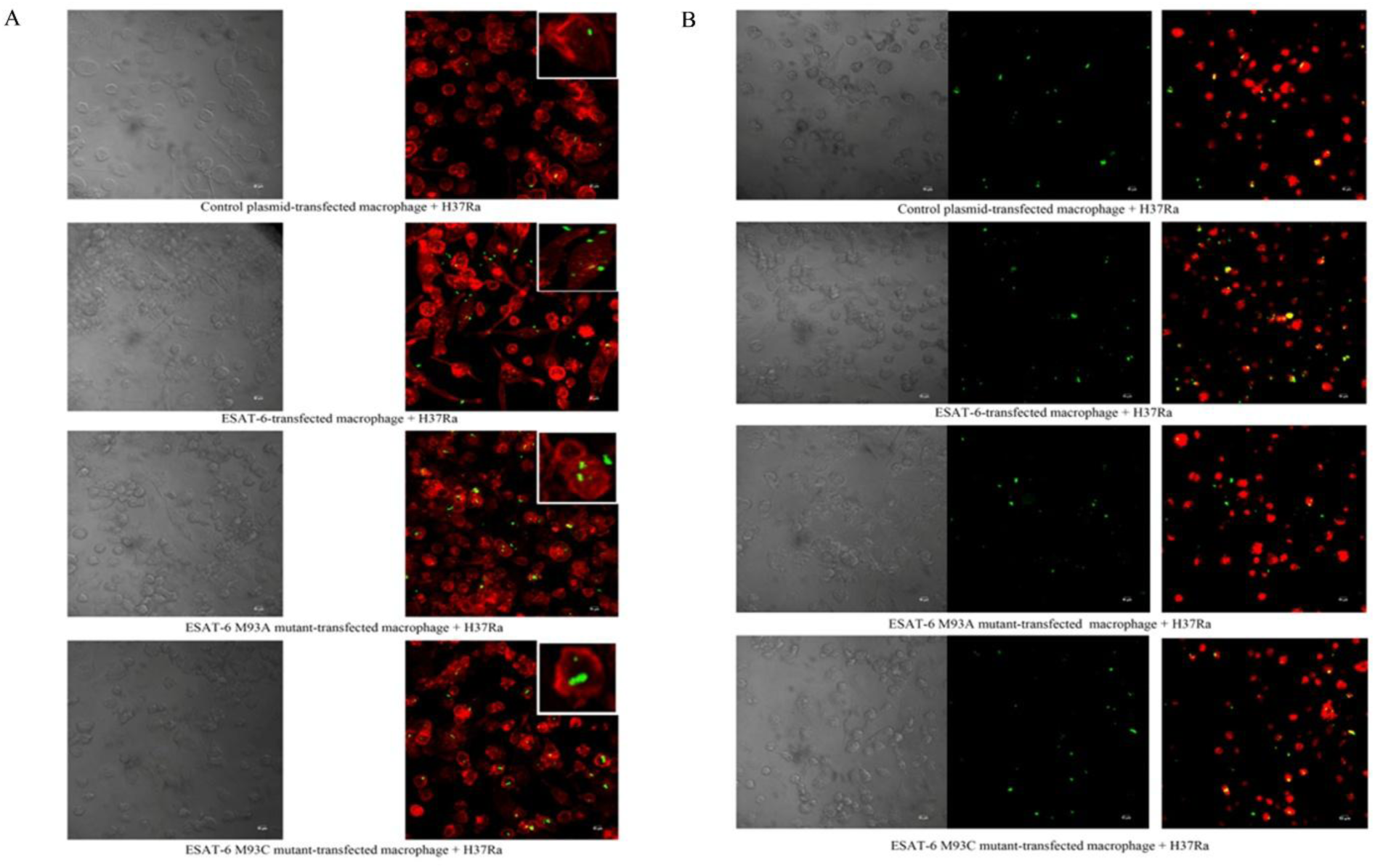
ESAT-6 increases the F-actin content, transforms the macrophages to spindle shaped and reduces phagosome acidification. These effects are partially reversed by its mutants. **A)** Phalloidine red staining was used to visualize F-actin in H37Ra (GFP expressing, green) - infected macrophages expressing ESAT-6 or its mutants, 24h post infection. **B)** Lysotracker Red staining was used to probe phagosome acidification in H37Ra (GFP expressing, green) - infected macrophages expressing ESAT-6 or its mutants, 24h post infection. The images are representatives of 3 independent experiments.

Further, we did *in silico* modeling to check for any structural changes caused by the M93 mutation in ESAT-6 and no considerable change was visible (Fig. 9). Along with this, the interaction of cytoplasmic methionine sulfoxide reductase (MsrB) of macrophage with ESAT-6 was predicted.

**Figure 9.**
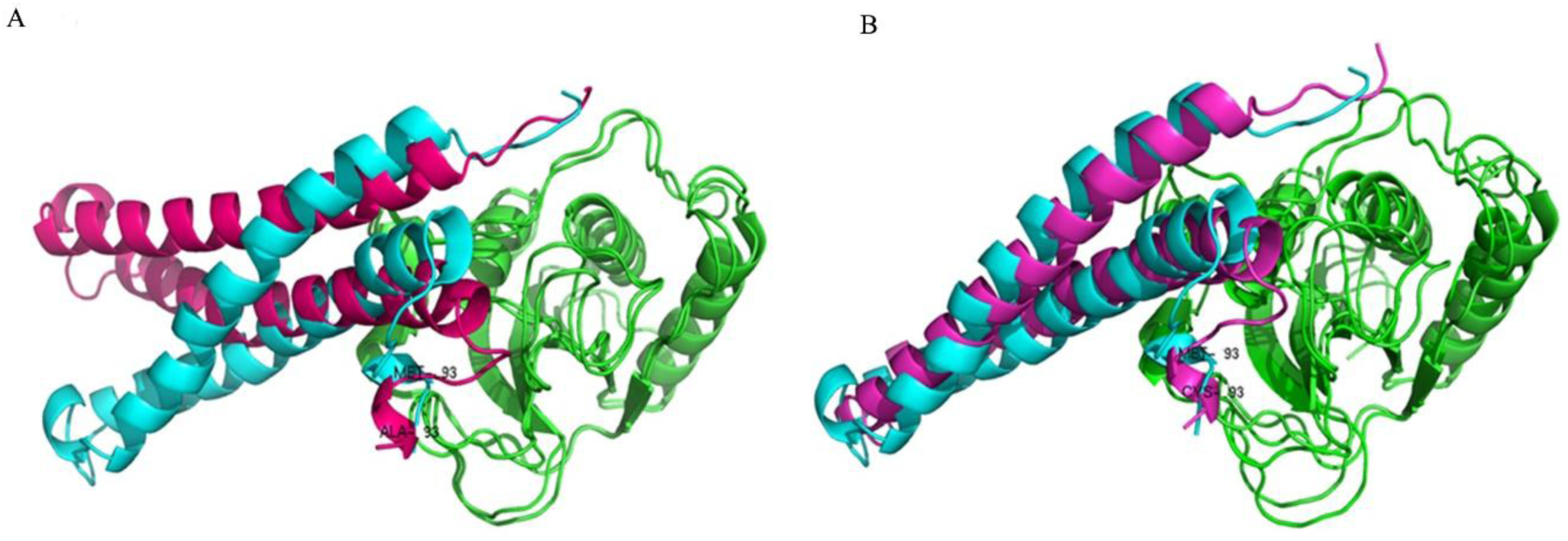
*In silico* modeling predicts that M93 mutations do not affect the structure of ESAT-6 considerably and the C-terminal flexible arm of ESAT-6 interacts with methionine sulfoxide reductase. **A)** ESAT-6 and ESAT-6 M93A mutant interact with MsrB. **B)** ESAT-6 and ESAT-6 M93C mutant interact with MsrB. (blue=ESAT-6, pink=ESAT-6 mutant and green=MsrB).

## 4. Discussion

Our study was rooted on the differential expression of cofilin1 of macrophages in an *in vitro* infection experiment. We guessed that the downregulation of cofilin1 and the upregulation of phosphocofilin1 in macrophages by virulent *M. tuberculosis* leads to enrichment of filamentous actin which in turn leads to improper phagosome processing. Increased phagosome acidification in infected macrophages expressing constitutively active cofilin1 strongly supports this assumption. We also showed that sorafenib which activates cofilin1 resulted in increased killing of *M. tuberculosis* in infected macrophages, giving an indication on the potential use of this kinase inhibitor for a host modulation therapy against tuberculosis.

Since cofilin1 is positively regulated by ROS we wanted to determine whether there is any effect by the ROS downregulating secreted protein ESAT-6 of *M. tuberculosis*, on cofilin1 or not. We found that ESAT-6 downregulates cofilin1 and results in decreased phagosome acidification. ESAT-6 and ESAT-6 like proteins are prominently secreted proteins of *M. tuberculosis*[28,29]. ESAT-6 is secreted from the RD1 genomic region of *M. tuberculosis* and it is absent in vaccine strain BCG[30]. Also, a mutation in the PhoP regulator contributes to the impaired secretion of ESAT-6 and attenuation of the avirulent *M. tuberculosis* strain, H37Ra[31]. In this study we propose a mechanism for downregulation of ROS by ESAT-6. Even though we do not provide clear evidence for the direct scavenging of ROS by methionines of ESAT-6, we could show that mutation of M93 methionine can abrogate ROS resistance imparted by the protein. The peculiar structural features of ESAT-6 lead us to the above assumption as mentioned earlier and we also suggest that the interactions of ESAT-6 with host cell receptors/proteins may happen by the virtue of methionine aromatic motif (methionine and an aromatic amino acid alternate in this motif) like sequence in the C-terminal arm since there is a phenylalanine near to M93[32].

The proposed mechanism of action of ESAT-6 on the antioxidant function wherein its multiple methionines contributing to its virulence makes us to speculate that similar mechanism may hold good in the case of ESAT-6 like proteins, for example, CFP-10. Since the *M. tuberculosis* genome contains many duplications of the Region of Difference (RD)[33] and most of the ESAT-6 like proteins, PE-PPE proteins and other hypothetical proteins which are secreted abundantly, but do not appear to have a role in the growth of the bacterium outside the host, a concerted action of all of these proteins may contribute to the virulence of the pathogen. The mechanism we proposed also implicates the possible scavenging of ROS by different biomolecules like carotenoids and sugars like mannose in various pathogens, might be contributing to their virulence.

Even though the RD1 region, secreting ESAT-6 and CFP-10 is absent, the vaccine strain BCG also contains ESAT-6 like proteins[34] and it is a fact that BCG is ineffective in adults. It is known that inefficient phagosome maturation and antigen presentation prevents ideal and long-term memory of T-cells[35]. Therefore, we think knocking out of abundantly secreted ESAT-6 like proteins from BCG genome may increase the efficiency of the vaccine.

## Author Contributions

Conceptualization, P.P.M and S.M; methodology, P.P.M, S.M, R.J.R and S.K.C; software, P.P.M and S.K.C.; validation, P.P.M, S.M, R.J.R and S.K.C; formal analysis, P.P.M and S.M; investigation, P.P.M, S.M and R.J.R; resources, S.M; data curation, P.P.M and S.M; writing—original draft preparation, P.P.M and S.M; writing—review and editing, P.P.M and S.M; visualization, P.P.M and S.K.C; supervision, S.M; project administration, S.M; funding acquisition, S.M. All authors have read and agreed to the published version of the manuscript.

## Acknowledgments

The authors thank Dr. Ajay Kumar, Dr. Jackson James, Dr. Omkumar, Dr. V.V Asha and Dr. Suparna Sen Gupta for providing some of the reagents.

